# Antibiotic induced biofilm formations in *Pseudomonas aeruginosa* strains KPW.1-S1 and HRW.1-S3 are associated with increased production of eDNA and exoproteins, increased ROS generation, and increased cell surface hydrophobicity

**DOI:** 10.1101/2022.07.19.500676

**Authors:** Abhinash Kumar, Saurav K Saha, Paromita Banerjee, Tapas K Sengupta

## Abstract

*Pseudomonas aeruginosa* is a medically important bacteria due to its ability to form biofilm and is also an opportunistic pathogen. *Pseudomonas aeruginosa* has the intrinsic ability to form biofilm as one of the defense mechanisms for their survival. The fact that it can form biofilms on various medical implants makes it more harmful clinically. Although various antibiotics are used to treat *Pseudomonas aeruginosa* infections, previous studies have shown that sub-MIC levels of antibiotics cause biofilm formation in this type of bacteria. The present study thus deals with the effect of the aminoglycoside antibiotic gentamicin on the biofilm dynamics of two *Pseudomonas aeruginosa* strains KPW.1-S1 and HRW.1-S3. Biofilm formation was seen to be increasing with increased gentamicin concentrations in growth media. Confocal laser scanning microscopy and scanning electron microscopy accompanied with other biochemical tests deduced that biofilm-forming components like exoproteins, eDNA, and exolipids as exopolymeric substances in *Pseudomonas aeruginosa* biofilms were increased in the presence of gentamicin. An increase in reactive oxygen species generation along with increased cell surface hydrophobicity was also seen in both strains when treated with gentamicin. The observed increase in the adherence of the cells accompanied by an increase in exopolymeric substances, eDNA, and exolipids may have largely contributed to the increased biofilm production by the *Pseudomonas aeruginosa* strains KPW.1-S1 and HRW.1-S3 under the stress of the antibiotic treatment.

## Introduction

*Pseudomonas aeruginosa* is a major opportunistic pathogen that causes diseases in immune-compromised individuals and patients with cystic fibrosis and thermal injuries (Joffe et al., 1992), thereby has become the model organism for Gram-negative biofilm. The potential of these bacteria is to form biofilms which lead to an efficient way for them to survive in the presence of antibiotics (Hall-Stoodley et al., 2004). Due to the formation of biofilms, they can be highly resistant up to 1000 times more than their planktonic brethren counterparts (Hoyle& Costerton, 1991). About 65% of infectious bacterial diseases in humans have been found to be caused by biofilms (Potera, 1999). Most medical devices are susceptible to colonization by pathogenic bacteria, and these biofilms are often sources of chronic infections (Schembri et al., 2002). With the progress of medical sciences, lots of medical devices are being used in human treatment which in turn is leading to increased opportunities for biofilm forming bacteria such as within intravenous catheters (Tran & Tran, 2012), urinary catheters (Donlan, 2002), cardiac pacemakers (Santos et al., 2011), as well as in chronic wound infections (Malic et al., 2013; Percival et al., 2012). In chronic bacterial infections, bacteria form a biofilm which is very difficult to eradicate. Different facets provide the resistance of antimicrobial factors, sorption to or reaction with the various components of EPS antibiotics converted into sublethal doses (Singh et al., 2010) slow-growing cells in the biofilms (Borriello et al., 2004), presence of phenotypically different “persister” cells (Spoering& Lewis, 2001; Suci& Tyler, 2003).

Antibiotics are natural products that are used in antimicrobial warfare and hence, antibiotics are considered to be “miracle drugs” used to treat many bacterial diseases in humans as well as in animals (Shlaes& Bradford, 2018). Although bacterial biofilms can be employed for bio remedial purposes (Dasgupta et al., 2013, 2015), they are found to be resistant to biocides, some antibiotics, and the immune responses of the host. Biofilms are considered alarming due to their resistant nature to antibiotics. Numerous studies have depicted that the sub-inhibitory concentration of antibiotics can induce bacterial biofilm formation and thus provide resistance to the bacteria (Penesyan et al., 2020). Biofilm is used as a resistant mechanism by the bacteria upon treatment with antibiotics. A proposed bacteriolysis effect of gentamicin explains this antibiotic’s efficacy (Wang et al., 2019). This antibiotic has the unique property of inhibiting bacterial protein synthesis. Thus, biofilm formation causes a reservoir of problems in the treatment of recurrent or lethal infections in humans and animals (Costerton et al., 1999).

The microorganisms comprising the biofilm live in self-made extracellular polymeric substances (EPS) that form their nearby environment, involving mainly polysaccharides, proteins, nucleic acids, and few lipids (Costa et al., 2018). The signal from the environment, such as alterations in pH, temperature, and the presence of inhibitory molecules like antibiotics, trigger the expression of a cascade of genes in bacteria (Fang et al., 2016), resulting in the secretion of extracellular polymeric substances (EPS) and bacteria form biofilm by masking cellular three-dimensional network within the EPS (Donlan, 2002). Thus, the microbes in biofilm live in a self-made matrix of EPS that creates their nearby environment. EPS are mainly composed of polysaccharides, proteins, nucleic acids, and lipids (Branda et al., 2005). They provide mechanical stability and minimize the effect of toxic chemicals on bacterial cells within a biofilm. The basic architectural units of biofilms are micro-colonies including different communities of microbial cells embedded into the extra polymeric substance matrix. Microcolonies are made up of 10–25% of cells and 79– 90% of the EPS matrix (Costerton et al., 1999).

EPS matrix defends biofilm cells from different unfavorable environmental situations, such as ultraviolet radiation, and sudden changes in pH values (H. C. Flemming et al., 2007). There are channels between microcolonies, through which water flows. These water channels work in biofilm as a modest circulatory system dispensing nutrients to all microcolonies and receiving the detrimental metabolites (Costerton and Lewandowski, 1995). Extracellular DNA (eDNA) is also a significant component of the matrix in *Pseudomonas. aeruginosa* biofilm, where it functions as a connector of cells (Yang et al., 2007). Bacteria are continuously evolving; thereby incorporating new ways to evade the effects of antibacterial drugs and hence, infections are getting more critical. The potential to create biofilm is one of the predominant ways through which bacteria develop resistance and get accustomed to antibiotics. Thus, bacterial biofilm causes chronic diseases as they display increased antibiotic adaptation and defend against phagocytosis and possibly other constituents of the adaptive as well as innate immune systems of a host (Biofilms and Their Role in Pathogenesis | British Society for Immunology). Reactive oxygen species (ROS) play an essential role in the host’s innate immune response to microbial pathogens. ROS has recently become one of the fascinating signaling molecules in higher eukaryotes, however, the role of ROS in regulating microbial physiology and structure is yet to be fully discovered. When *P. aeruginosa* strains were exposed to sublethal concentration of polymyxin B, increased oxidative cell burst was observed. (Lima et al., 2019) suggested that polymyxin B increases ROS production via the quinone-NADH oxidoreductase pathway; as a result, the oxidative burst produces free radicals which are responsible for damaging lipids, and DNA resulting in cell death. In a study by (Stephanie Da Cruz Nizer et al., 2021), it was seen that cell death along with hydroxyl radical production was caused in the presence of antibiotics by promoting the Fenton reaction. In the natural environment, microorganisms prefer to live as aggregates rather than planktonic cells which is an important parameter of forming aggregates or adhesion is cell surface hydrophobicity. It is significant that with the addition of disinfectants, glutaraldehyde, and chlorhexidine, some percentage of *Pseudomonas sp* cells moved to the hydrophobic phase (Fitzgerald et al., 1992; Mcdonnell & Russell, 1999). Various factors affect hydrophobicity like temperature and growth conditions. When some *Pseudomonas* isolates were grown at 37°C, cell surface hydrophobicity got shifted to being hydrophilic whereas cells grown at a relatively lower temperature showed hydrophobic behavior (Hori et al., 2009).

It is therefore important to study how sub-MIC concentrations of antibiotics regulate biofilm formation in *Pseudomonas aeruginosa*. Thus, the aim of the present study was to delineate the effect of gentamicin which is an aminoglycoside antibiotic on the biofilm formation by two environmental isolates of *Pseudomonas aeruginosa* KPW.1-S1 and HRW.1-S3. We investigated the biofilm morphology, production of exopolymeric substances, ROS generations and changes in cell-surface hydrophobicity of the strains due to the effect of sub-MIC doses of gentamicin.

## Materials and Methods

### Bacterial strains and growth conditions

The bacterial strain, **KPW.1-S1** and **HRW.1-S3** isolated from Kolkata port water and Haldia river water respectively and identified as *Pseudomonas aeruginosa* (Dasgupta et al., 2013) are used for the present study. The environmental isolates **KPW.1-S1** and **HRW.1-S3** were deposited at MTCC and numbered MTCC 10087 and MTCC 10088 respectively. The strains were grown in Bushnell Haas media with 2% added glucose. For that, 10 ml of BH 2% glucose media was inoculated with the strains individually and kept in shaking conditions overnight at 37° C.

### Determination of sub-minimum inhibitory concentrations of Gentamicin

Seed cultures of *Pseudomonas aeruginosa* KPW.1-S1 and HRW.1-S3 were obtained by inoculating in 10 ml of BH media containing 2 % glucose at 37°C for 10 -12 hours. Post growth, *Pseudomonas aeruginosa* KPW.1-S1 and HRW.1-S3 were adjusted in such a way that the starter culture should have 3×10^6^ cells (CFU) /ml in BH media (2% glucose). 3 ml of culture was taken in a 15 ml test tube with different concentrations of antibiotics (viz. 0.25, 0.5, 0.75, 1, 1.25, 1.5, 2, 2.5, 5, and 10 µg/ml of gentamicin sulphate) and incubated at 150 rpm, 37°C for 18 hours. The next day, OD was measured at 600nm in Schimadzu Spectrophotometer.

### Biofilm Load Assay

Biofilm load assays were performed on 24 well polystyrene coated plates (NUNC). Three independent experiments were performed for the quantification of biofilm load. A fixed number of cells (∼ 3×10^6^ cells/ml) of KPW.1-S1 and HRW.1-S3 in BH 2% glucose were grown on plates in the absence or presence of desired concentrations of antibiotics for 24 hours at 37° C at 150 rpm. After the incubation period, the planktonic cells were discarded, and the wells were then washed with PBS (Phosphate Buffer Saline) twice, 1.5 ml 0.1% CV (crystal Violet) solution was added to each well for staining, followed by incubation for 45 minutes, at room temperature. CV was removed from each well and washed using PBS (three times). 24 well microtiter plate was inverted on tissue papers to remove any excess liquid. The plate was allowed to dry. 1 ml of 30% acetic acid solution (Zegans et al., 2009) was added and then kept at shaking condition for 30 minutes at 37° C, 150 rpm. The O.D. was measured at λ=600nm (Merritt et al., 2005) on a 24 well cum 96 well plate reader (BioTekELx 800). The average O.D. determined by the ELISA plate reader for the control wells was subtracted from the O.D. of all test wells.

### Determination of Colony Forming Unit (CFU) of bacteria present in the biofilm matrix

*Pseudomonas aeruginosa* KPW.1-S1 and HRW.1-S3, grown in 24 well plates were maintained at 3×10^6^ cells/ml, in the presence or absence of antibiotics (0.75 and 1.5 µg/ml of gentamicin) in BH 2% glucose. The plates were then incubated for 24 hours at 37°C under shaking conditions (150 rpm). The planktonic cells were then removed carefully, and the wells were washed with PBS (Phosphate Buffer Saline) two times. Post-washing, 1 ml of PBS was added to each well. The biofilm matrix was disrupted by gentle pipetting followed by serial dilution. 100 µl suspension of it was spread on nutrient agar plates. The plates were then incubated at 37°C overnight. The next day, bacterial colonies were counted in different dilution factors.

### Confocal Laser Scanning Microscopy (CLSM)

*Pseudomonas aeruginosa*, KPW.1-S1, and HRW.1-S3 biofilm load assay in the presence and absence of antibiotic (Gentamycin) were assessed by the Confocal Laser Scanning Microscope (CLSM) image sacks (Allan et al., 2002). The growth of both the strains was performed on glass coverslips in BH (2% glucose) media (24 hrs at 37° C and 150 rpm) in 24 well plates. After 24 hrs, the coverslips were washed with PBS twice and stained with 0.005% acridine orange (w/v) solution in the dark for 5 minutes and then washed with PBS (thrice) and mounted with 5% glycerol (v/v) on a clean glass slide. The edges of the coverslips were sealed with transparent nail polish. The glass slides were viewed under a Carl Zeiss CLSM-710 microscope utilising a 488 nm excitation wavelength and emission was observed in the range of 293-586 nm. Topological parameters (Thickness, Volume, Skewness, Kurtosis) of biofilm were calculated using the software provided.

### Scanning Electron Microscopy (SEM)

To get a better resolution of *Pseudomonas* biofilms (cell and EPS levels) and an understanding of the cellular changes, Scanning Electron Microscopy was performed (Xie et al., 2013). The strains, KPW.1-S1 and HRW.1-S3 were grown on glass coverslips in the presence or absence of antibiotics (0.75 and 1.5 µg/ml Gentamicin) for 24 hrs at 37° C, 150 rpm in 24 well plates. Coverslips were kept in 2.5% Glutaraldehyde for 2 hrs and then washed with PBS thrice for 15 min, followed by the addition of 0.1% Osmium tetraoxide and incubation in the dark for 40 mins. Then they were washed with PBS thrice. After this, the cells were washed by alcohol gradation of 30%, 50%, 70%, 90%, and 100% every 10 mins. The cover slip specimen was allowed to dry in a vacuum and then was viewed under a Carl Zeiss microscope using Smart software.

### Visualization of the presence of eDNA in biofilms by fluorescence microscopy

Extracellular (eDNA) and intracellular DNA (iDNA) of *Pseudomonas aeruginosa* (KPW.1-S1 and HRW.1-S3) biofilms grown in the presence or absence of antibiotics (0.75 and 1.5 µg/ml gentamicin) were visualized by dual staining using DAPI (0.5 µg/ml in PBS) and EtBr (1. 5 µg/ml in PBS) under a fluorescence microscope (ZEISS, ApoTome.2). The KPW.1-S1 and HRW.1-S3 strains were grown on glass coverslips in BH 2% glucose media for 24 hours at 37° C under shaking conditions (150 rpm). After incubation, the coverslips were washed with PBS twice and then coverslips were put in EtBr (1.5 µg/ml in PBS) for 5 minutes. The coverslips were then washed with PBS, and then incubated with DAPI (0.5 µg/ml in PBS) for 5 minutes, then again washed with PBS. Coverslips were put on the glass slides covered with 5 % glycerol. These samples were then observed under a fluorescence microscope. An excitation wavelength of 353 nm, 558 nm, and emission wavelength of 465 nm and 575 nm were used for DAPI and EtBr respectively.

### Quantification of eDNA in biofilm by spectrophotometry

*Pseudomonas aeruginosa* biofilms (KPW.1-S1 and HRW.1-S3) strains were grown in the presence or absence of antibiotics 0.75 and 1.5 µg/ml of gentamicin in BH 2% glucose media for 24 hours at 37° C under shaking condition (150 rpm) in 24 well plates. Post-incubation, the planktonic cells were removed carefully, and the wells were washed with PBS (Phosphate Buffer Saline) two times. 1 ml of TE buffer was added and biofilms were scraped from the surface (gentle pipetting) (Nakao et al., 2012). The suspension containing the scrapped biofilms was centrifuged at 11,000 RPM for 2 min and then the supernatants were transferred to a new tube and the remaining bacterial cells were removed by filtering through a 0.22 µm filter (Millipore, Ireland). The extracellular DNA (eDNA) was estimated using EtBr (1 µg/ml). 30 µl of both the filtered supernatant and EtBr were mixed well. 50 µl of the mixture was used for observation under a fluorescence microscope fluorescence. (Bonasera et al., 2007). Anexcitation wavelength of 254 nm and emission wavelength of 616 nm were used respectively.

### Visualization of extracellular lipid by fluorescence microscopy

*Pseudomonas aeruginosa* biofilms (KPW.1-S1 and HRW.1-S3) strains were grown in the presence or absence of antibiotics (0.75 and 1.5 µg/ml gentamicin) on glass coverslips in BH 2% glucose media for 24 hrs at 37° C under shaking condition 150 rpm in 24 well plates. After 24 hours of incubation, the coverslips were washed with PBS twice and stained with Nile red (2 µg/ml in PBS) in dark for 10 minutes and then washed with PBS and mounted with 5% glycerol (v/v) on a clean glass slide. The edges of the coverslips were sealed with transparent nail polish. The glass slides were observed under a fluorescence microscope (ZEISS, ApoTome.2). The wavelengths used were 530 nm for excitation and 580 nm for emission.

### Estimation of extracellular lipids by spectrophotometry

*Pseudomonas aeruginosa* biofilms (KPW.1-S1 and HRW.1-S3) were grown in BH 2% glucose for 24 hours at 37° C under shaking conditions (150 rpm) in 24 well plates. After that, biofilms were washed with PBS twice and 0.9 % NaCl was added. Biofilms were then scraped from the surface using gentle pipetting (Nakao et al., 2012). After centrifugation (11,000 RPM for 2 min), the supernatant was collected in a new tube and the remaining bacterial cells were removed through a 0.22 µm filter (Millipore, Ireland). The extracellular lipids of biofilms were quantified by using Nile red (Chen et al., 2009). Briefly, 5 µl of supernatant of biofilms were mixed with 292 µl of 20% DMSO and 3 µl of Nile Red (0.5 µg/ml in 75% Glycerol). After that, it was heated at 40° C for 10 minutes and the fluorescence was measured using 530 nm as excitation wavelength and emission wavelength was detected at 580 nm.

### Extracellular Protein analysis

*Pseudomonas aeruginosa* biofilms (KPW.1-S1 and HRW.1-S3) were grown in BH 2% glucose for 24 hours at 37° C under shaking conditions (150 rpm) in 24 well plates. After incubation, careful removal of the planktonic culture was done. Biofilms were then washed with PBS twice. 1 ml of 0.9% NaCl was added and biofilms were scraped from the surface using gentle pipetting (Nakao et al., 2012). The scrapped biofilms were centrifuged at 11000 rpm for 2 minutes. After centrifugation, the supernatants were transferred to a new tube and the remaining bacterial cells were removed by filtering through a 0.22 µm filter (Millipore, Ireland). The extracellular protein concentrations were measured using the Bradford method. In brief, 200 µl of supernatant of biofilm and 50 µl of Bradford dye reagent were mixed. After 5 minutes, 200 µl of the mixture was used to measure the absorbance at 595 nm with the help of a microplate reader (Wu & Xi, 2009).

### Extracellular polysaccharide analysis

*Pseudomonas aeruginosa* biofilms (KPW.1-S1 and HRW.1-S3) strains were grown in the presence or absence of antibiotics (0.75 and 1.5 µg/ml gentamicin) on glass coverslips in BH 2% glucose media 24 hrs at 37° C under shaking condition 150 rpm in 24 well plates. The planktonic culture was carefully removed after incubation. The biofilms were then washed with PBS twice. 1 ml of 0.9% NaCl was added and biofilms were scraped from the surface using gentle pipetting (Nakao et al., 2012). The scrapped biofilms were centrifuged at 11000 rpm for 2 minutes. After centrifugation, the supernatants were transferred to a new tube and the remaining bacterial cells were removed via a 0.22 µm filter (Millipore, Ireland). The Anthrone method was used for extracellular polysaccharides determination. In brief, 80 µl of the supernatant was mixed with 160 µl of Anthrone reagent (0.125% Anthrone in 94.5 % H_2_SO_4_). Samples were heated in a water bath at 100° C for 14 minutes and then cooled at 4°C for 5 minutes. The absorbance at 625 nm was measured using a microplate reader (BioTeK) (Wu & Xi, 2009).

### Qualitative analysis of Reactive Oxygen Species (ROS) by Fluorescence microscopy

Reactive Oxygen Species (ROS) were measured using Dichloro-dihydro-fluorescein diacetate (DCFH-DA), an oxidation-sensitive fluorescent probe. *Pseudomonas* aeruginosa KPW1.S1 and HRW.1-S3 were grown in 24 well plates, and biofilms were washed with PBS (twice). Biofilm was treated with DCFH-DA (10 µM) for 30 min at 37°C. The fluorescence intensity of DCF was measured using a fluorescence spectrophotometer at an excitation wavelength of 485 nm and an emission wavelength of 535 nm. The sample for the negative control was treated with 5% H_2_O_2_ followed by the DCFDA Treatment for 30 min. The samples were washed with 1 ml of saline and resuspended in 100 µL of it .10µl is loaded onto the slides and visualized using an epifluorescence microscope (Olympus –IX equipped with Hamamatsu camera) with a blue excitation filter (460-495nm) and emission filter (510-550 nm).

### Microbial Adhesion to the Hydrocarbon (MATH Assay)

*Pseudomonas aeruginosa* (KPW.1-S1 and HRW.1-S3) strains were grown under different conditions to measure the hydrophobicity using the MATH assay. It was performed using hydrophobic hydrocarbon n-Hexadecane. The optical density of bacterial cells was measured at 600 nm. The cell density was then adjusted to 0.6 by dissolving in PBS. 1.5ml of adjusted cell suspension was mixed with 0.5ml of n-hexadecane. The mixture was then vortexed for 30 seconds. The separated aqueous layer was collected and its O.D. was measured at 600 nm (A_1_). The surface hydrophobicity was assessed by the formula: [1-(A_1_/A_0_)]*100.

### Statistical analysis

GraphPad Prism 6 was used to plot and analyze the data. All the analyses were carried out in triplicates and stated as mean ± standard error. Two-tailed Student’s t-test was used to determine significant differences between means. Statistically significant differences were indicated by asterisks (*P<0.05; **P<0.01; ***P<0.001)

## Results

### Sub-minimum inhibitory concentrations of Gentamicin

The minimum inhibitory concentration of gentamicin against KPW.1-S1 and HRW.1-S3 is 2 µg/ml (**Figure 1)**. For both the strains, *Pseudomonas aeruginosa* HRW.1-S3 and HRW.1-S3 (**Figure 1 a** and **1 b**), MIC of gentamicin was also found to be 2 µg/ml. The growth of both strains was inversely proportional to the concentration of gentamicin. For further studies of the biofilm of both strains, the gentamicin concentrations are considered to be 0.75 µg/ml and 1.5 µg/ml.

**Figure 1.**
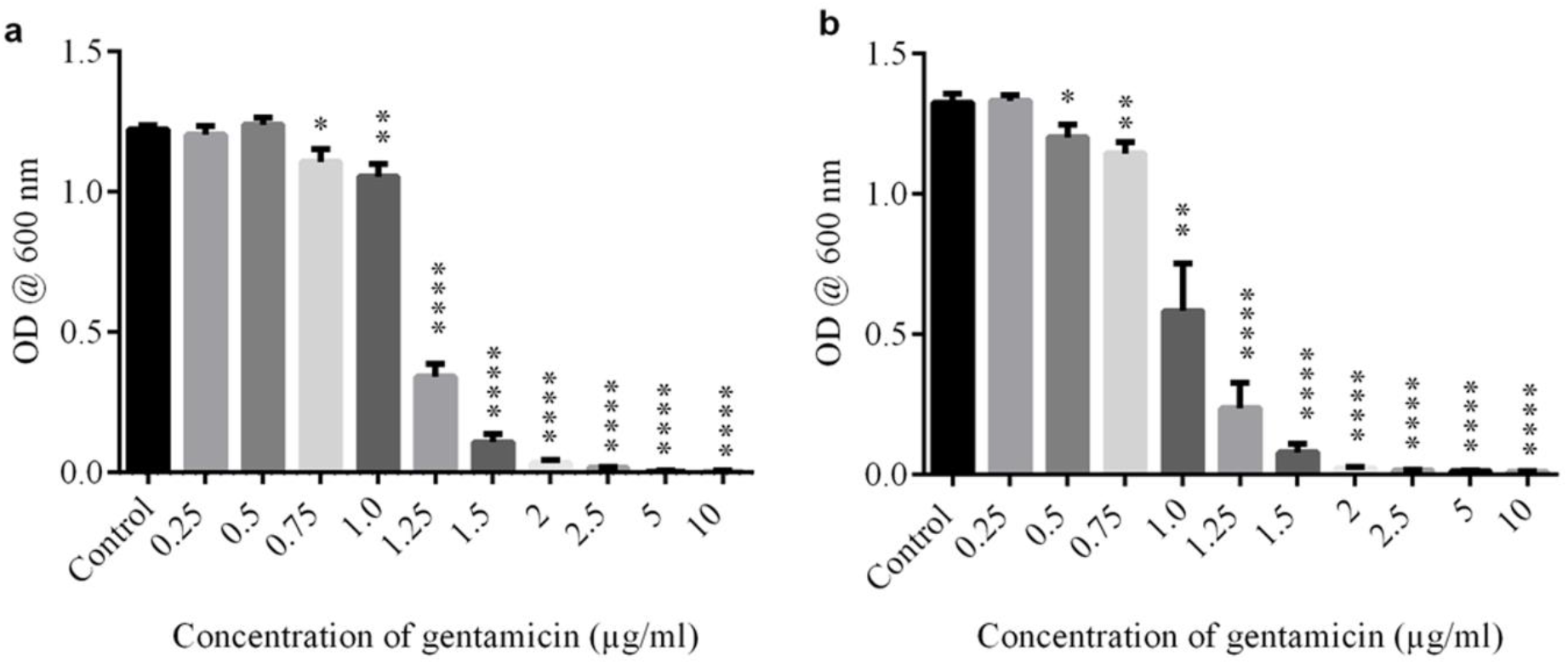
Minimum Inhibitory Concentration (MIC) of *Pseudomonas aeruginosa* KPW.1-S1 **(a)** and HRW.1-S3 **(b)**. with different concentrations of antibiotics (viz. 0.25, 0.5, 0.75, 1, 1.25, 1.5, 2, 2.5, 5 and 10 µg/ml of gentamicin sulphate) The data represent mean values ± SE of N=6. Statistically significant differences are indicated by asterisks when compared to control (\**P* < 0.05; \*\**P* < 0.01; \*\*\**P* < 0.001; \*\*\*\**P* < 0.0001 two-tailed Student’s *t-test*).

### Effect of Gentamicin on biofilm formation of *Pseudomonas aeruginosa* KPW.1-S1 and HRW.1-S3

The biofilm load assay was done for both strains of *Pseudomonas aeruginosa* KPW.1-S1 and HRW.1-S3 grown in BH-2 % glucose in the absence and presence of 0.75 and 1.5 µg/ml of gentamicin. The biofilm load of the strain KPW.1-S1 was compared with control (without gentamicin). Biofilm load was found to be increased significantly at 1.5 µg/ml of gentamicin for KPW.1-S1 **(Figure 2 a)** compared to control. Whereas in HRW.1-S3, biofilm load was found to be increased in both the gentamicin concentrations (0.75 and 1.5 µg/ml) when compared to control **(Figure 2 c)**. The CV staining indicates that biofilms were formed on the bottom as well as the wall of 24 well plates **(Figure 2 b** and **Figure 2 d)**. It was observed that for *Pseudomonas aeruginosa* KPW.1-S1 and HRW.1-S3, the presence of sub-inhibitory concentrations of the gentamicin could induce a significant increase in biofilm loads.

**Figure 2.**
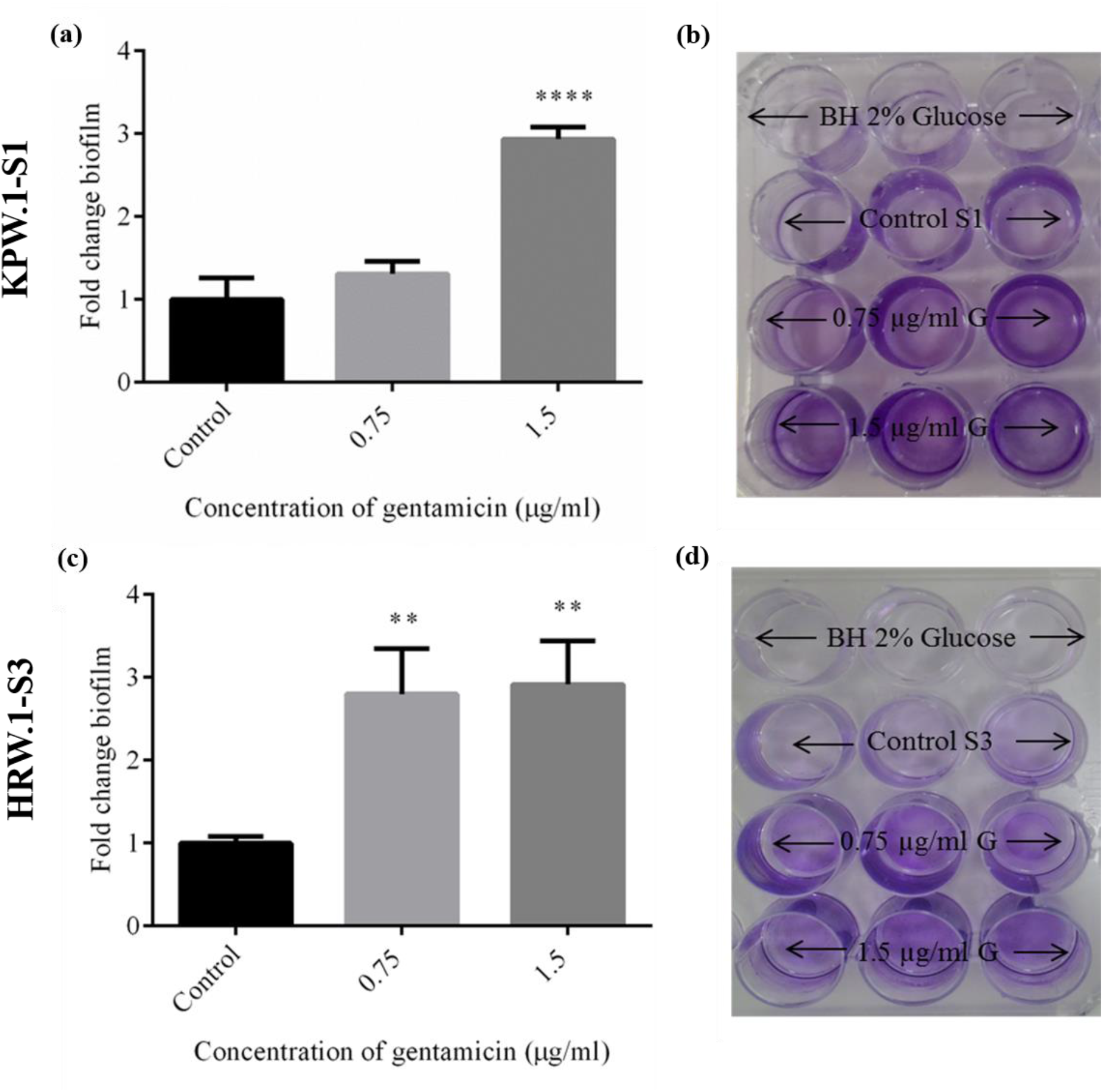
Biofilm load assay of *Pseudomonas aeruginosa* KPW.1-S1 and HRW.1-S3in presence of **(a)** and **(c)**0.75 µg/ml, and 1.5 µg/ml of gentamicin and **(b)** and **(d)** 1% CV staining of biofilm plate. The data represent mean values ± SE of N=6. Statistically significant differences are indicated by asterisks when compared to control (\**P* < 0.05; \*\**P* < 0.01; \*\*\**P* < 0.001; \*\*\*\**P* < 0.0001 two-tailed Student’s *t-test*).

### Topological parameters and architecture of KPW.1-S1 and HRW.1-S3 biofilm

Biofilm architecture was visualized and different topographical parameters were measured using confocal laser scanning microscopy (CLSM) of *Pseudomonas aeruginosa* strains KPW.1-S1 and HRW.1-S3 were grown in BH-2 % glucose in the absence and presence of 0.75 and 1.5 µg/ml of gentamicin. There was increased biofilm production observed when the KPW.1-S1 strain was grown in the presence of gentamicin **(Figure 3 b** and **Figure 3 c)** as compared to control **(Figure 3 a)**. The average biofilm thickness of KPW.1-S1 was found to be 7.02 μm and 5.68 μm in the presence of 0.75 and 1.5 µg/ml of gentamicin respectively, whereas it was found to be 3.84 μm in the case of the control (**Figure 4 a)**.

**Figure 3.**
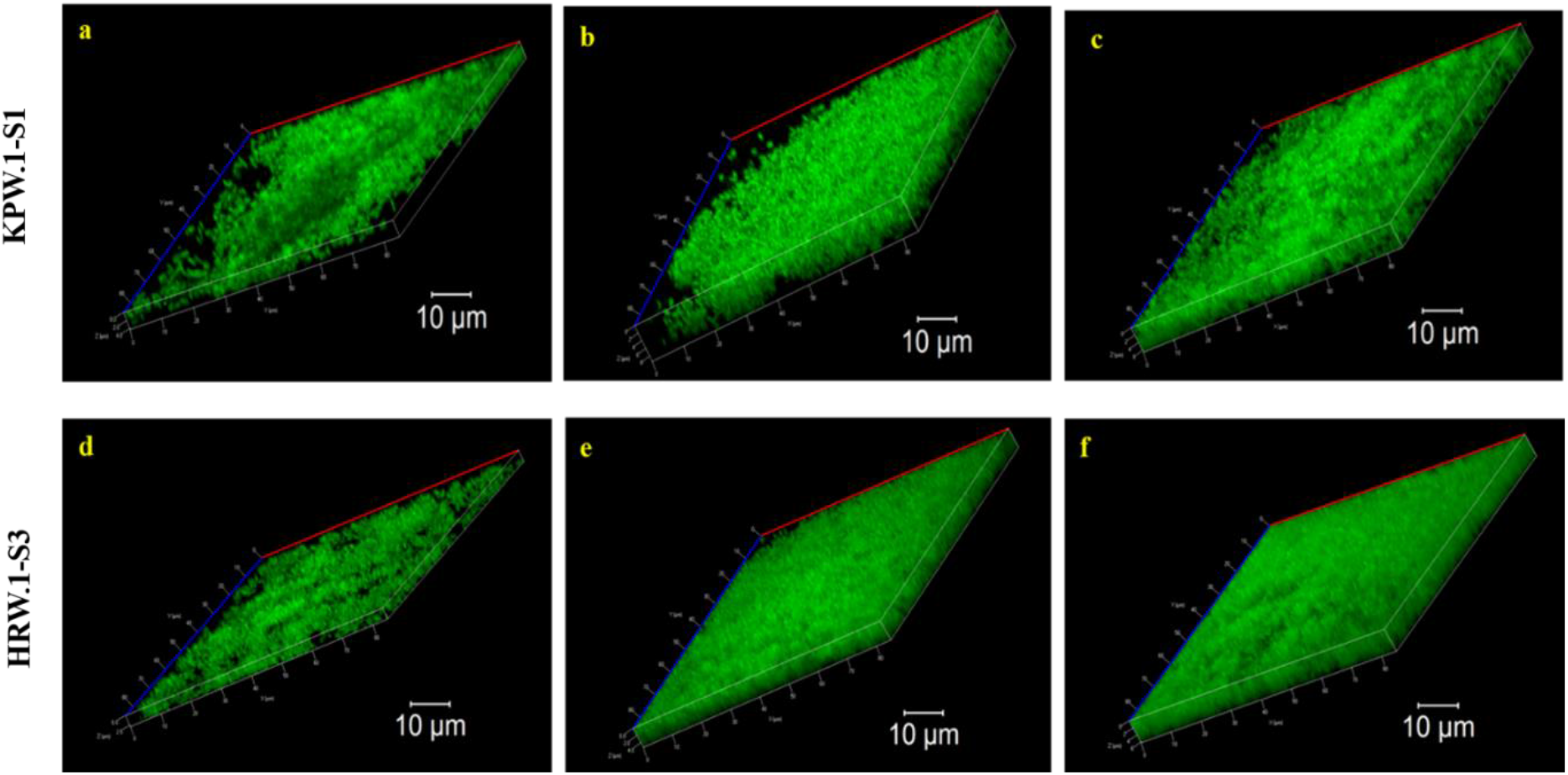
Confocal laser scanning microscopy image of biofilmof*Pseudomonas aeruginosa*, KPW.1-S1 and HRW.1-S3 in the presence and absence of Gentamycin. Representative images of KPW.1-S1, **(a)** control, **(b)** 0.75 µg/ml, and **(c)** 1.5 µg/ml and HRW.1-S3, **(d)** control, **(e)** 0.75µg/ml, and **(f)** 1.5 µg/ml of gentamicin. Bars, 10 *μ*m.

**Figure 4.**
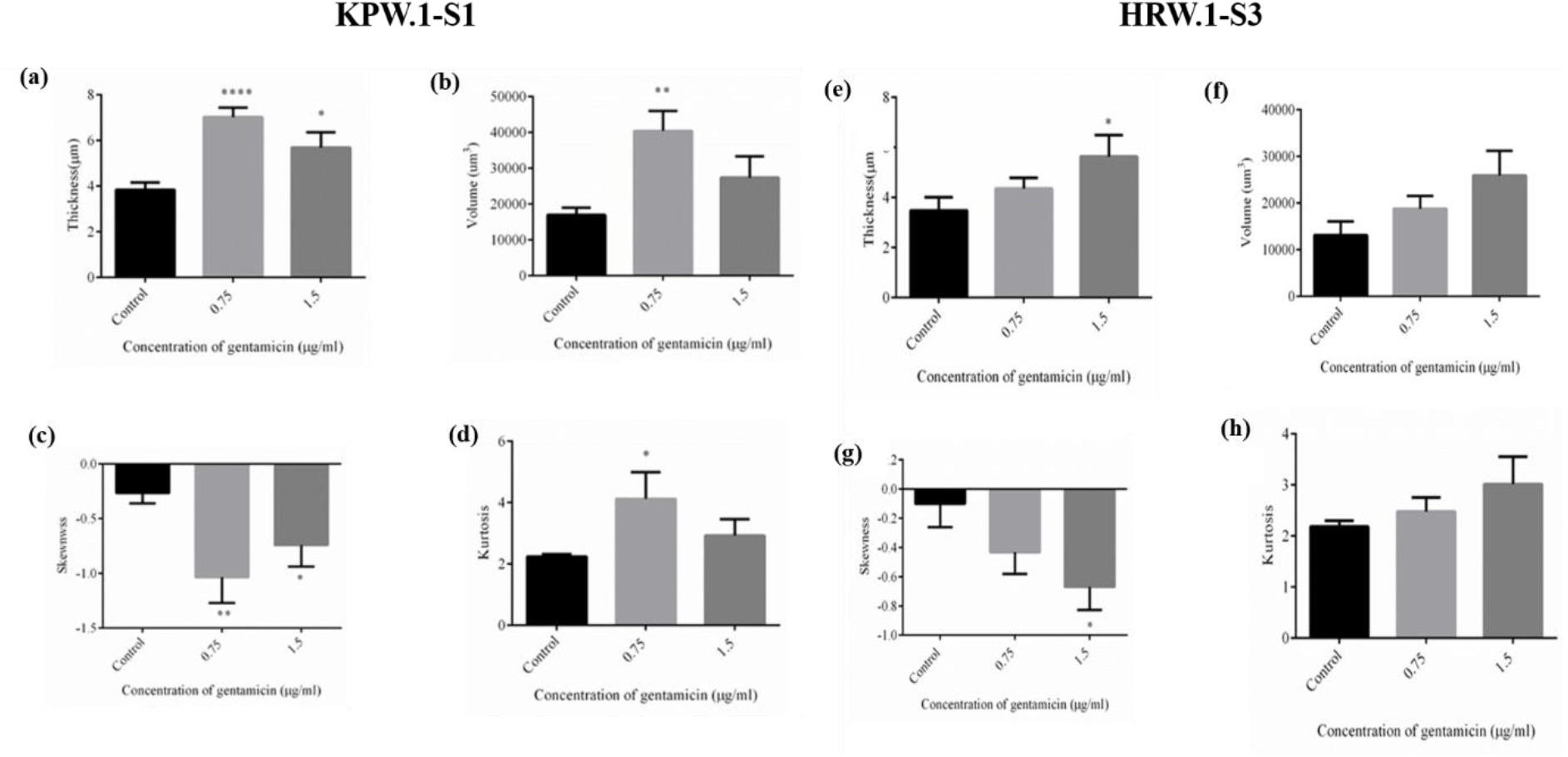
Topological features of biofilms formed by *Pseudomonas aeruginosa* KPW.1-S1and HRW.1-S3 in presence of 0.75 µg/ml, and 1.5 µg/ml of Gentamicin after 24 h of incubation: (**a) and (e)** Thickness, (**b)and (f)** Volume, (**c)and (g)** Skewness and (**d) and (h)** Kurtosis. The data represent mean values ± SE of N=8. Statistically significant differences are indicated by asterisks when compared to control (\**P* < 0.05; \*\**P* < 0.01; \*\*\**P* < 0.001; \*\*\*\**P* < 0.0001 two-tailed Student’s *t-test*).

In the case of *Pseudomonas aeruginosa* HRW.1-S3, the biofilm production was enhanced compared to control **(Figure 3 d)** in the presence of gentamicin **(Figure 3 e** and **Figure 3 f)**. The average thickness of HRW.1-S3 was found to be 4.356 and 5.637 μm in the presence of 0.75 and 1.5 µg/ml gentamicin respectively, whereas it was found to be 3.485 μm in the case of the control **(Figure 4 e)**. The thickness, biovolume, and kurtosis were also found to be increased as compared to control while the Skewness decreased. Overall, in the presence of gentamicin, for both the strains KPW.1-S1 and HRW.1-S3, thickness, biovolume, skewness, and kurtosis of biofilms were found to be changed as compared to the control conditions **(Figure 4 a-h)**. Also, the mean thickness **(Figure 4 a.e)** and volume **(Figure 4 b.f)** for both the strains were found to be inversely proportional to skewness **(Figure 4 c,g)** in the presence of gentamicin. Since, higher skewness indicates a lack of porosity, negative values of skewness **(Figure 4 c,g)** indicate that the surface is dominated by holes or valleys and hence has greater porosity which enables the biofilm to gain more nutrients and other important factors which results in the greater thickness of the biofilm. The thickness, biovolume, and kurtosis were significantly increased while the Skewness which signifies the presence of void spaces within a biofilm decreased significantly as compared to the control.

Biofilms formed by the isolated *Pseudomonas aeruginosa* KPW.1-S1 and HRW.1-S3 in the presence of 0.75µg/ml and 1.5µg/ml of gentamicin were analyzed by scanning electron microscopy (SEM). Microscopy images demonstrated that the distribution of bacterial micro-colonies was not homogenous throughout the glass cover surface in *Pseudomonas aeruginosa* KPW.1-S1 and HRW.1-S3 **(Figure 5 a-f)**. In some areas, extracellular material was observed connecting the bacteria. In contrast, control samples of both the strains showed that some of the bacterial cells adhere to the surface and very little extracellular matrix was present **(Figure 5 a,d)**. Biofilm load increased with increasing concentrations of gentamicin with a significant increase in EPS structure **(Figure 5 b,c,e,f)**. *P. aeruginosa* biofilms as seen for both strains under microscopy demonstrated an increase in biofilm density, as well as a significant increase in EPS with an increasing concentration of gentamicin, which was also observed.

**Figure 5.**
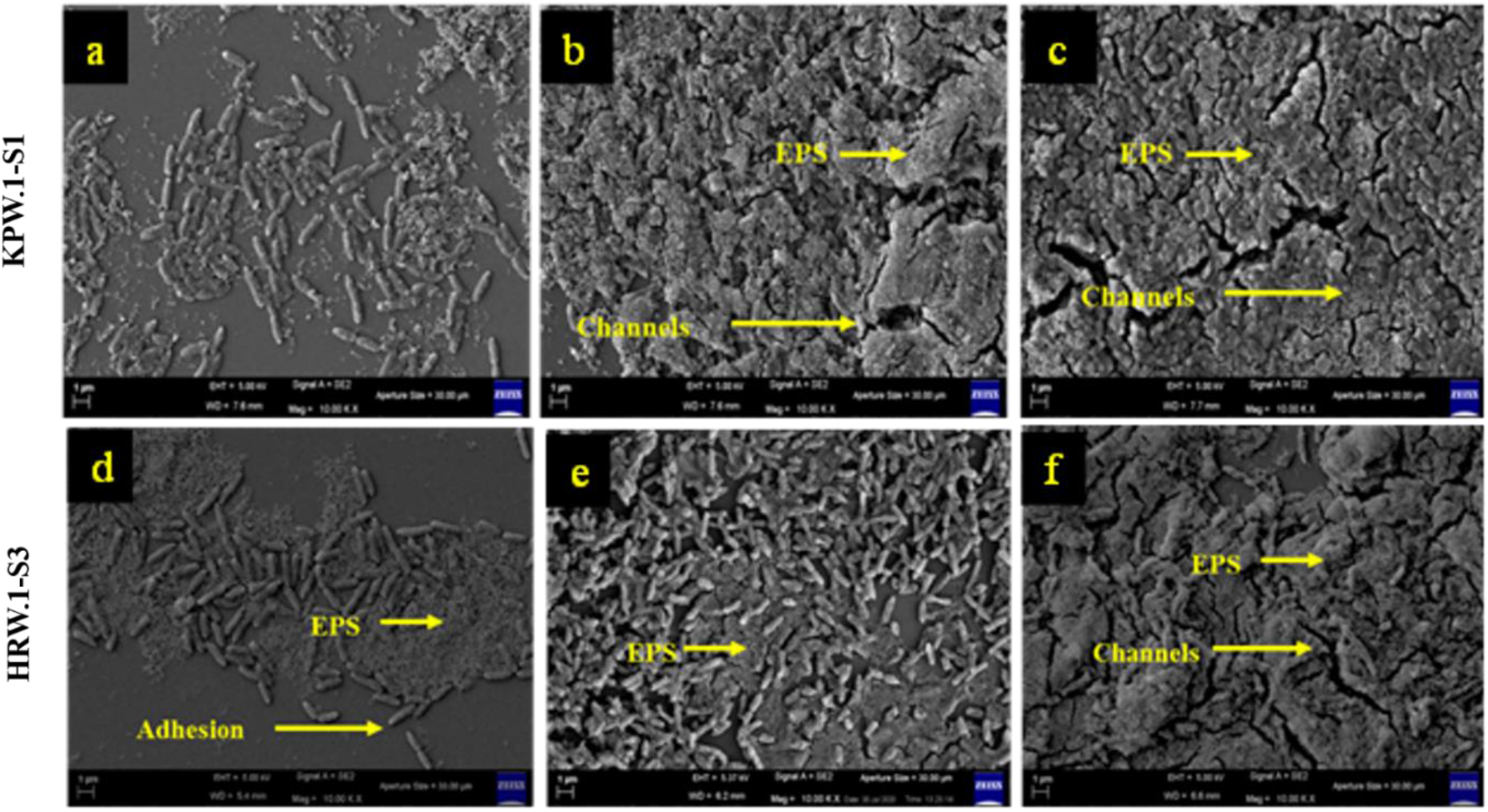
Representative images of Scanning Electron Micrograph of *Pseudomonas aeruginosa* KPW.1-S1**(a-c)** and HRW.1-S3**(d-f)**biofilm formed on the coverslips in presence of gentamicin **(a)**and **(d)**Control, **(b)** and **(e)** 0.75 µg/ml, **(c)** and **(f)** 1.5 µg/ml. Bar 1 µm.

### Visualization of the presence of eDNA in biofilms by fluorescence microscopy

Extracellular DNA (eDNA) and intracellular DNA (iDNA) of *Pseudomonas aeruginosa* KPW.1-S1 and HRW.1-S3 biofilms grown in the presence or absence of 0.75 and 1.5 µg/ml gentamicin were visualized by the dual staining by DAPI (0.5 µg/ml in PBS) and EtBr (1. 5 µg/ml in PBS) by using a fluorescence microscope (ZEISS, ApoTome.2). The red fluorescence was visible due to EtBr binding with eDNA of biofilm and blue fluorescence light was visible due to the internal DNA of bacterial cells of biofilms.

The microscopic data showed the presence of eDNA in the biofilm matrix and EtBr was bound with eDNA (red fluorescence). It was observed in the case of KPW.1-S1 that eDNA was more abundant in biofilms grown in presence of higher doses of gentamicin (**Figure 6 a**). Similarly for HRW.1-S3 also, eDNA was found to be more abundant in biofilms grown in presence of higher doses of gentamicin (**Figure 6 c**).

**Figure 6.**
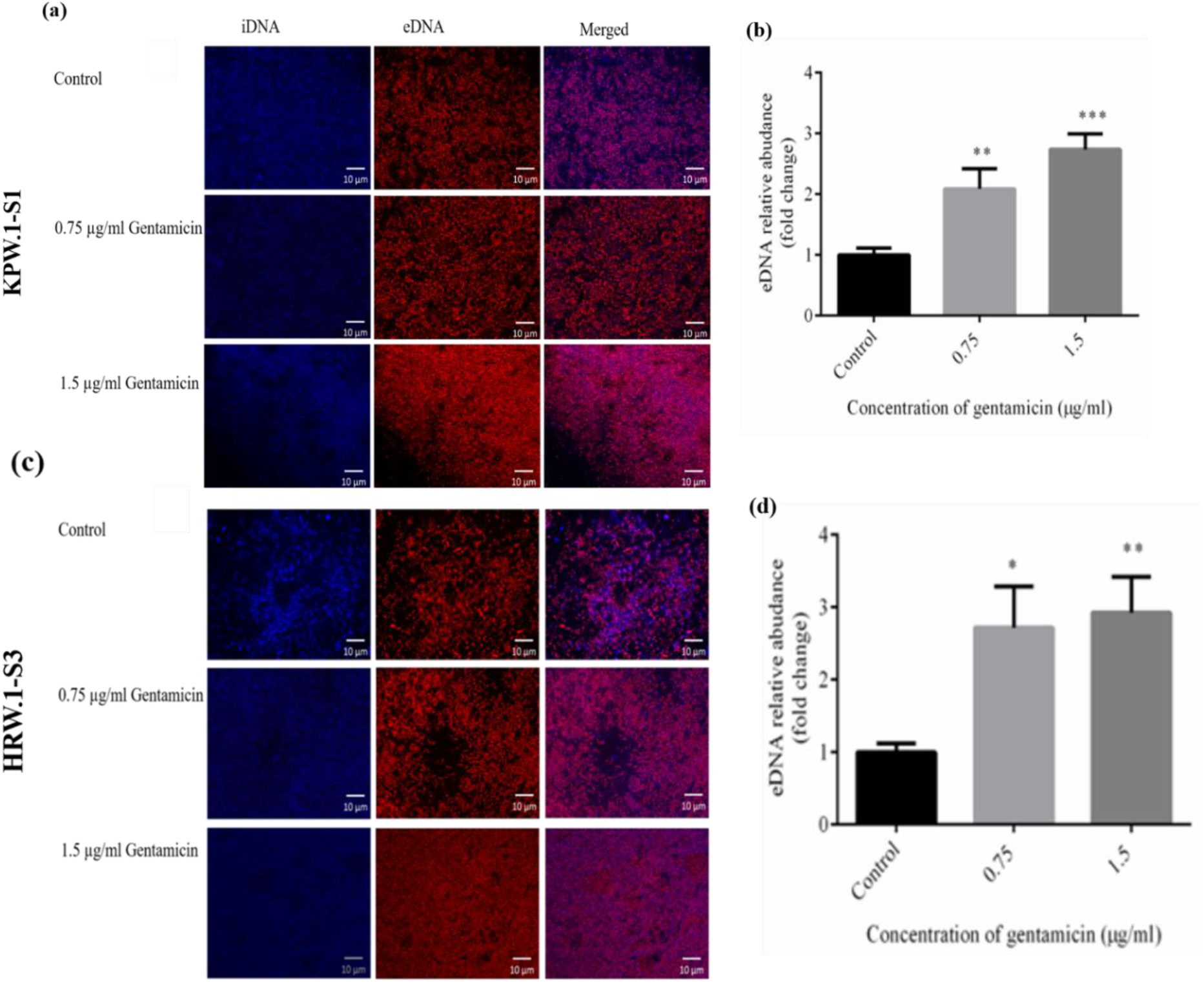
Representative images of Fluorescence Microscopy of Biofilm **(a)**KPW.1-S1 and **(c)** HRW.1-S3and (**b**) and **(d)** graph of eDNA in biofilm KPW.1-S1 and HRW.1-S3 respectively. The data represent mean values ± SE of N=9. Statistically significant differences are indicated by asterisks when compared to control (\**P* < 0.05; \*\**P* < 0.01; \*\*\**P* < 0.001; \*\*\*\**P* < 0.0001 two-tailed Student’s *t-test*).

The first studies on ethidium bromide (EtBr) for the quantification of nucleic acid were done in the mid-1960s. For decades, ethidium bromide (EtBr) has been the most often used fluorescent dye in nucleic acid gel electrophoresis. However, spectrofluorimetric-based DNA quantification has mostly been overlooked. For *Pseudomonas aeruginosa* KPW.1-S1 biofilms,2.09 and 2.73 folds rise in eDNA relative abundance were observed when the biofilms were allowed to form in presence of 0.75 µg/ml and 1.5 µg/ml of gentamicin, respectively compared to control condition (**Figure 6 b**).

For *Pseudomonas aeruginosa* HRW.1-S3, 2.71, and 2.92 folds relative increase in eDNA abundance was observed when biofilms were formed in the presence of 0.75 µg/ml and 1.5 µg/ml gentamicin respectively as compared to control (**Figure 6 d**). These results show that eDNA release contributes to some extent to gentamicin enhanced biofilm formation by *Pseudomonas aeruginosa* KPW.1-S1 and HRW.1-S3. The cumulative data indicate that gentamicin at sub-MIC induced eDNA release, mostly through cell lysis, thereby modifying the composition of the matrix, which might contribute to the observed increased biofilm formation.

### Quantification of Extracellular Protein

Analysis of exoproteins of *Pseudomonas aeruginosa* KPW.1-S1 biofilms indicated a 2.5 and 2.36 fold relative increase in exoprotein abundance when biofilms were subjected to 0.75 µg/ml and 1.5 µg/ml gentamicin respectively (**Figure 7 a**). *Pseudomonas aeruginosa* HRW.1-S3 biofilm exoproteins analyses showed a 2.48 and 1.8 fold relative increase in exoproteins abundance when biofilms were treated with 0.75 µg/ml and 1.5 µg/ml gentamicin respectively as compared to the control condition in absence of any antibiotic in eDNA abundance (**Figure 7 b**). These results indicate that exoproteins release contributes to a certain extent to gentamicin-induced biofilm formation by *Pseudomonas aeruginosa* KPW.1-S1 and HRW.1-S3.

**Figure 7.**
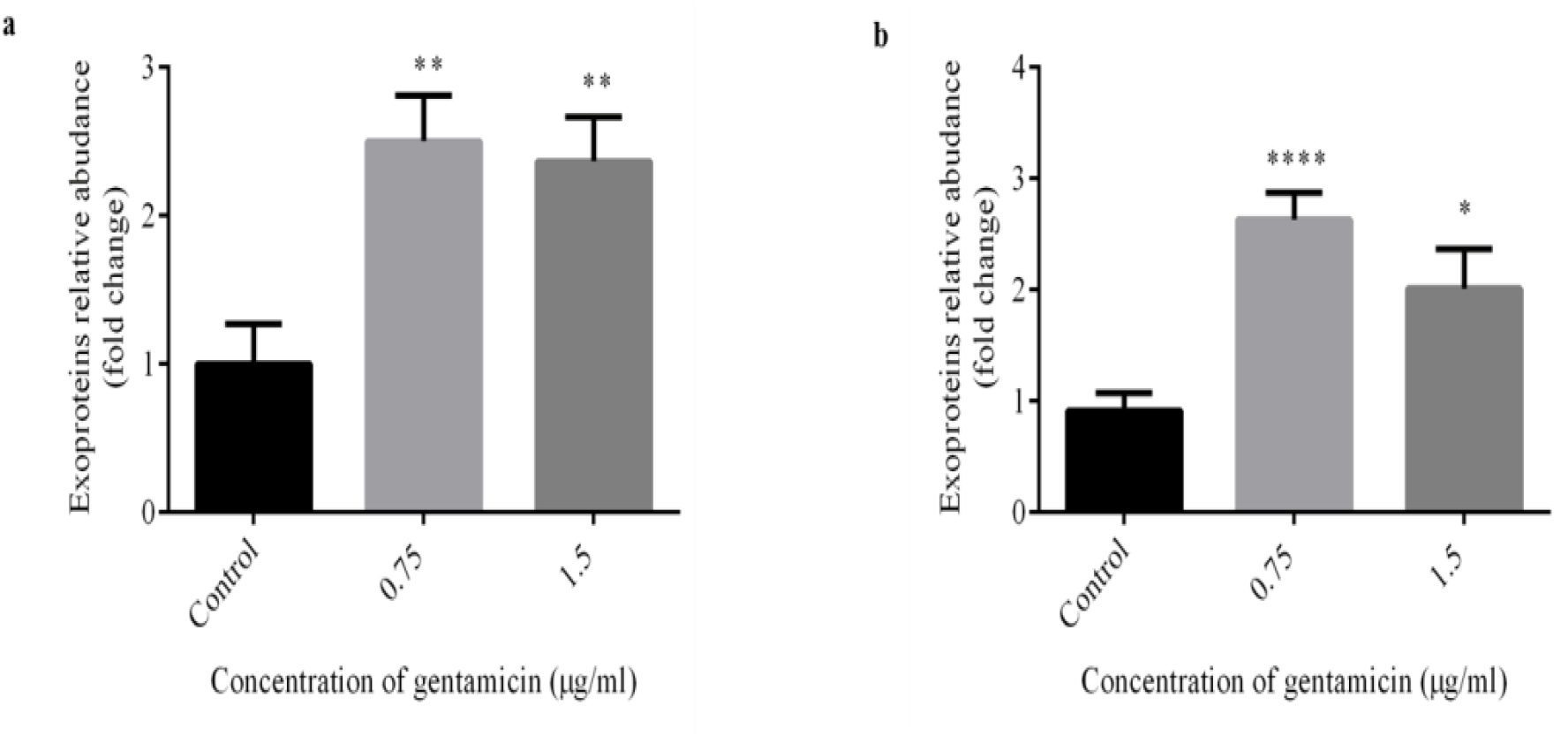
Estimation of exoproteins of *Pseudomonas aeruginosa* biofilm in presence of gentamicin **(a)**KPW.1-S1, and (b) HRW.1-S3. The data represent mean values ± SE of N=9. Statistically significant differences are indicated by asterisks when compared to control (\**P* < 0.05; \*\**P* < 0.01; \*\*\**P* < 0.001; \*\*\*\**P* < 0.0001 two-tailed Student’s *t-test*).

### Visualization and estimation of extracellular lipid

To visualize the relative abundance of lipid moieties in the biofilm structures, Nile red staining was done. Although the presence of lipids in EPS/Biofilm was observed in biofilms for all conditions, no significant alteration in distribution and abundance was visible in the microscopic images of Nile red-stained biofilms of *Pseudomonas aeruginosa* KPW.1-S1 and HRW.1-S3 grown in absence or presence of gentamicin as compared to control (**Figure 8 a**). Spectroscopic analysis of lipid content in *Pseudomonas aeruginosa* KPW.1-S1 biofilms revealed 1.39 and 1.80 fold relative increase in extracellular lipids abundance when biofilms were grown in presence of 0.75 µg/ml and 1.5 µg/ml gentamicin respectively when compared with the control (**Figure 8 b**).

**Figure 8.**
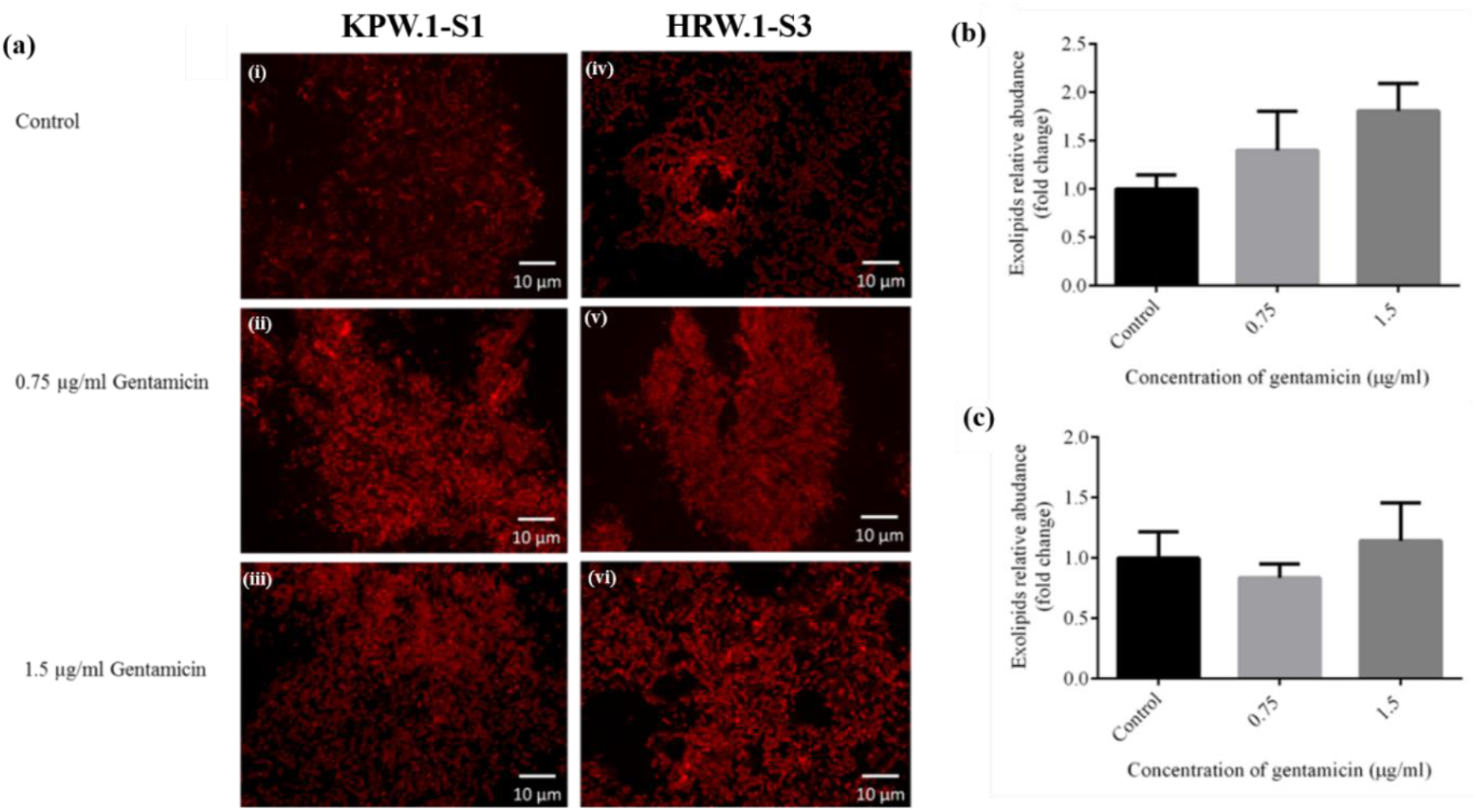
Extracellular lipids of biofilms were quantified using Nile red Nile red staining of biofilm **[a (i-iii)]** KPW.1-S1 and HRW.1-S3**[a (iv-vi)]** in Control **(i)** and **(iv)** and presence of antibiotic **(ii)**and **(v)**0.75 µg/ml, **(iii)** and (vi) 1.5 µg/ml.of gentamicin sulphate. Quantification of exolipids of KPW.1-S1 **(b)** and HRW.1-S3**(b)** is represented in the graph. The data represent mean values ± SE of N=3.

For *Pseudomonas aeruginosa* HRW.1-S3 biofilm, extracellular lipids were found to be reduced by 0.165 fold in presence of 0.75 µg/ml gentamicin while 1.5 µg/ml of gentamicin could induce 1.142 fold increase in relative abundance (**Figure 8 c**).

### Quantification of Extracellular Polysaccharides

Although the estimation of exopolysaccharides in *Pseudomonas aeruginosa* biofilms grown in the absence or presence of gentamicin showed differential abundance but the differences were found to be statistically insignificant.KPW.1-S1 biofilm exopolysaccharides analyses indicated 1.52 fold increase in the presence of 0.75 µg/ml gentamicin but almost no alteration was seen in presence of 1.5 µg/ml gentamicin (**Figure 9 a**). For *Pseudomonas aeruginosa* HRW.1-S3 biofilm, exopolysaccharide analyses revealed a 1.60-fold increase in presence of 0.75 µg/ml gentamicin while in the presence of 1.5 µg/ml of the antibiotic there was only a 0.042 fold reduction as compared to control (**Figure 9 b**).

**Figure 9.**
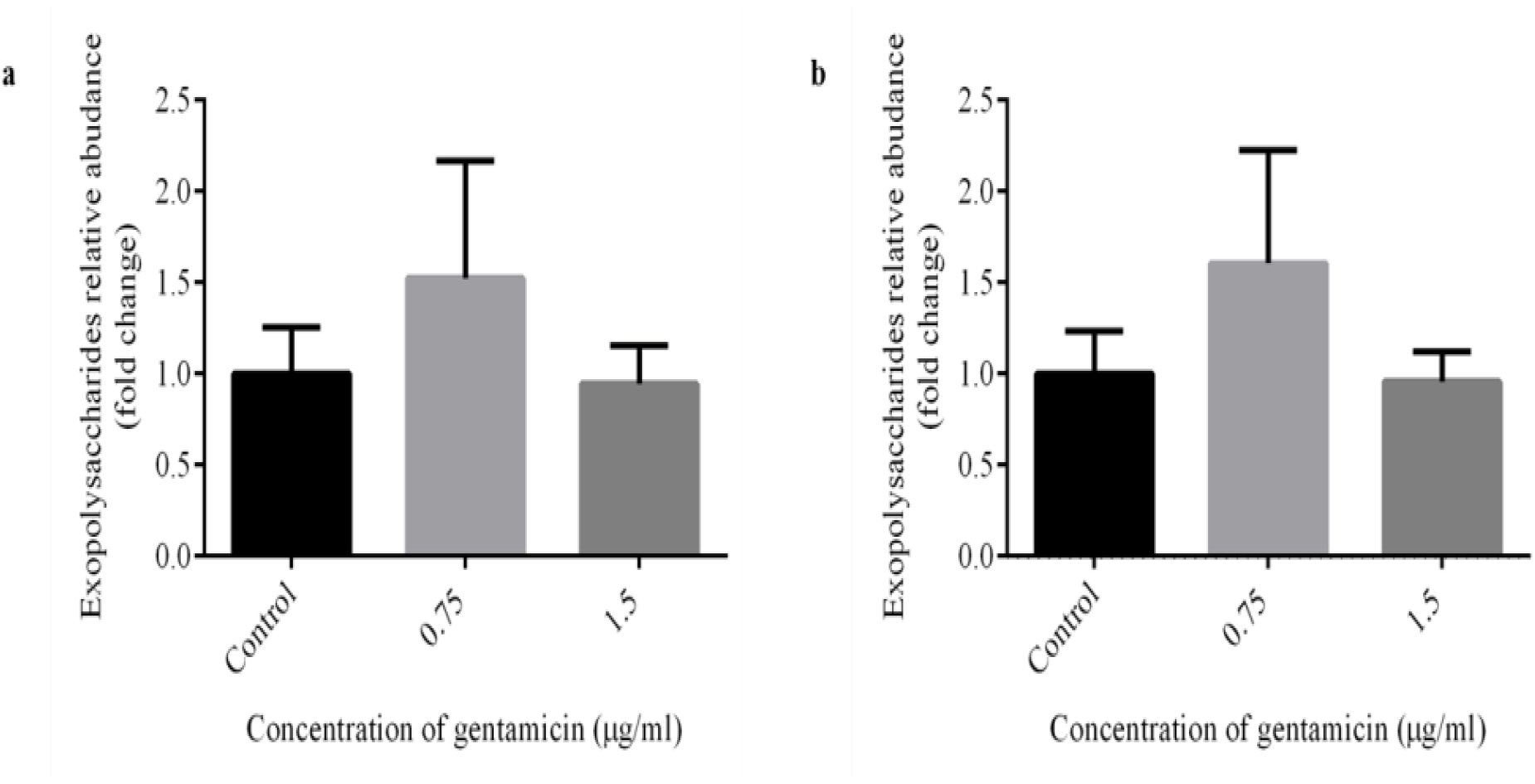
Extracellular polysaccharides quantification was performed using the Anthrone method of biofilm**(a)** KPW.1-S1and**(b)** HRW.1-S3. The data represent mean values ± SE of N=9 and N=8 respectively.

### Quantification of ROS generation in presence of gentamicin

The fluorescence microscopy indicated an increase in ROS production in both KPW.1-S1 and HRW.1-S3 in presence of gentamicin as compared to control (**Figure 10 a, 10 b**). The quantitative estimation of ROS production was done using the software GenBlue. In the case of KPW.1-S1, there was a 1.64 fold and 1.69 fold increase in ROS production compared to control cells in the presence of 0.75 µg/ml and 1.5 µg/ml gentamicin respectively (**Figure 10 c**). The HRW.1-S3 strains showed a 1.56- and 3.75-fold increase in ROS production in presence of 0.75 µg/ml and 1.5 µg/ml gentamicin respectively as compared to control (**Figure 10 d**).

**Figure 10.**
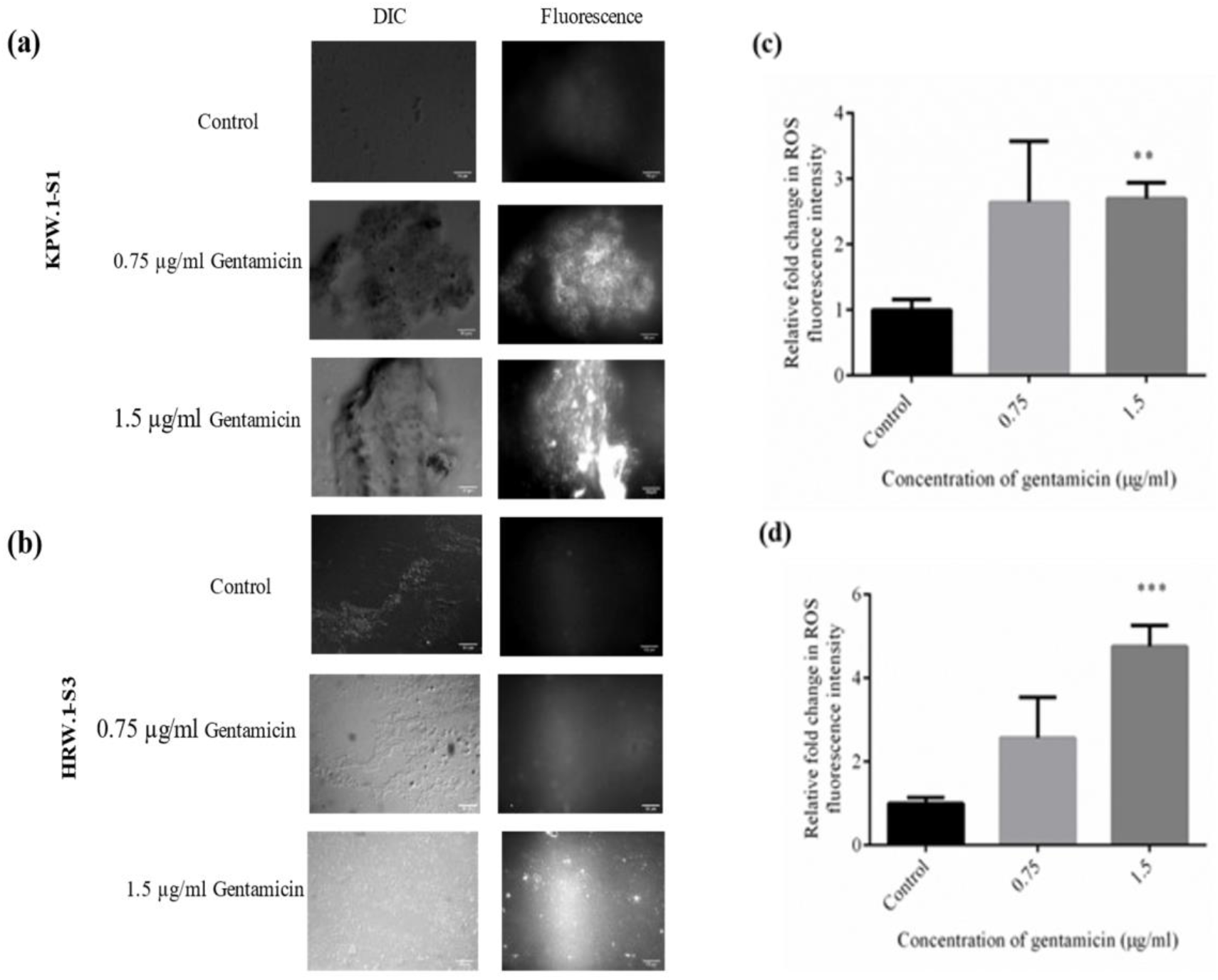
Amount of ROS accumulation in terms of fluorescence intensity of biofilm **(a)**KPW.1-S1 and **(a)**HRW.1-S3 in the presence of gentamicin. The intensity of ROS was quantified for biofilm **(a)**KPW.1-S1 and **(a)**HRW.1-S3. The data represent mean values ± SE of N=4. Statistically significant differences are indicated by asterisks when compared to control (\**P* < 0.05; \*\**P* < 0.01; \*\*\**P* < 0.001; \*\*\*\**P* < 0.0001 two-tailed Student’s *t-test*).

### Effect of gentamicin on bacterial adhesion properties

Cell surface hydrophobicity is an important factor depending on which bacteria form biofilms on different surfaces. MATH assay was used to assess cell surface hydrophobicity of both the strains in the presence and absence of gentamicin.

In the case of KPW.1-S1, the control cell was found to be on the slightly negative side of the hydropathy index with a value of -14 denoting a slightly hydrophilic nature. After the treatment with 0.75 and 1.5 µg/ml gentamicin respectively, the graphs shifted to the positive side with values of 3 and 6.366 respectively denoting a slightly hydrophobic nature (**Figure 11 a**).

**Figure 11.**
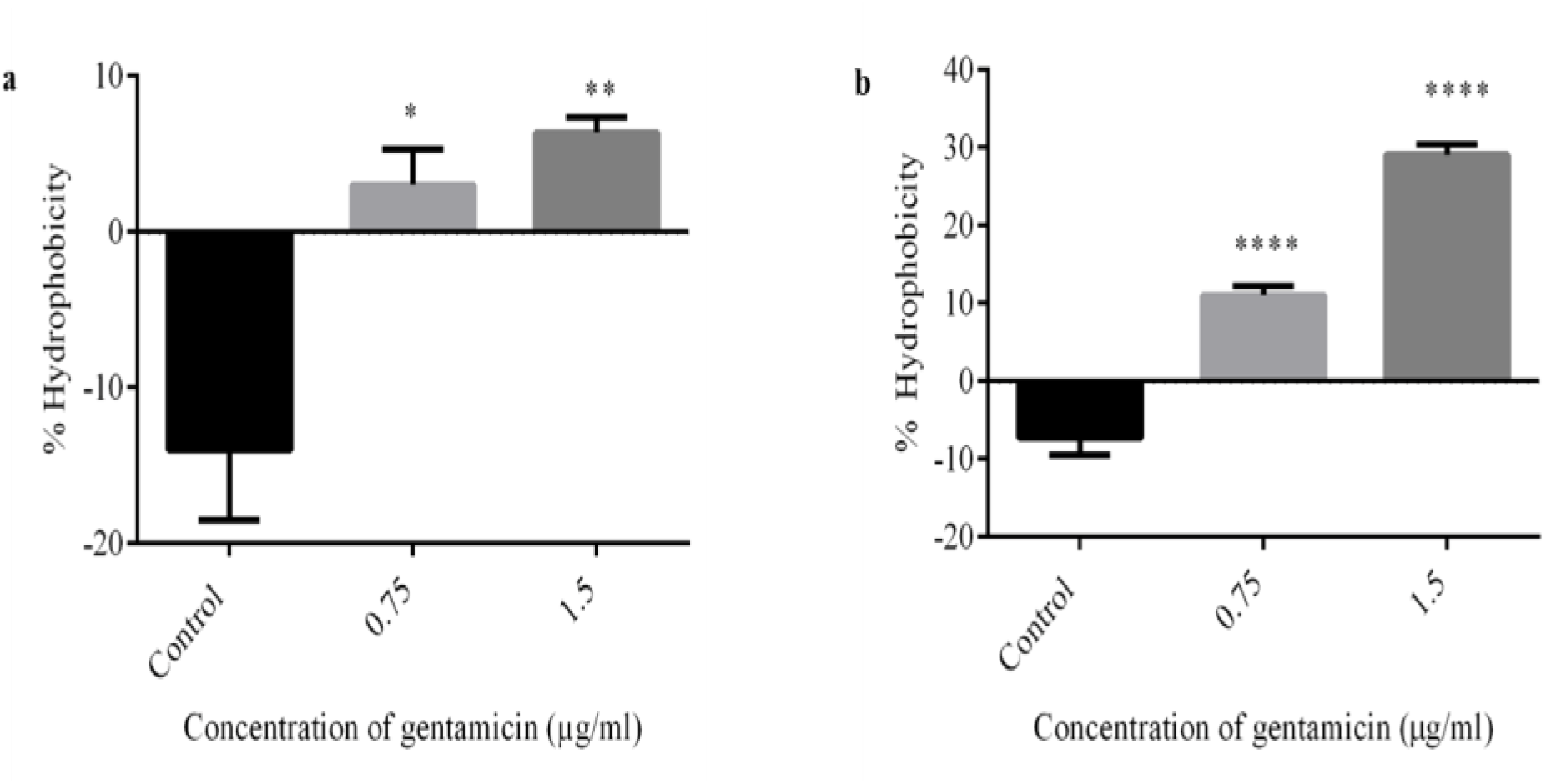
Estimation of adherence property of *Pseudomonas aeruginosa***(a)**KPW.1-S1 and (b) HRW.1-S3 in presence of gentamicin was done. Data represent mean ± SE of N=6. Statistically significant differences are indicated by asterisks when compared to control (\**P* < 0.05; \*\**P* < 0.01; \*\*\**P* < 0.001; \*\*\*\**P* < 0.0001 two-tailed Student’s *t-test*).

For HRW.1-S3, a similar trend was observed. The untreated cells (control) showed a negative score of -7.33 while after the addition of 0.75 and 1.5 µg/ml gentamicin the value shifted to the positive side with values of 11 and 29.06 respectively showing an increase in hydrophobicity (**Figure 11 b**).

## DISCUSSION

Biofilms have a critical role in causing and elevating chronic infections. Biofilm-forming bacteria have characteristics that distinguish them from planktonic ones, such as higher tolerance to antibiotics and biocide agents, the formation of different interactions, an increased rate of gene exchange, and phenotypic variant selection (Ciofu et al., 2017; H. C. Flemming& Wingender, 2010). Several bacteria isolated from hospital environments were found to be capable of surviving on different surfaces like catheters as biofilms, acting as a medium of infectious agents for hospitalized individuals as well as healthcare workers (Vincent, 2003). Interestingly, antibiotics at levels below the minimal inhibitory concentration (sub-MIC) can trigger the alteration in the formation of biofilm, virulence, and gene expression, which contributes to overall bacterial phenotypic and genetic resistance (Andersson & Hughes, 2014; Davies et al., 2006; Fajardo & Martínez, 2008; Song et al., 2016). In some clinical pathogens such as *Staphylococcus aureus, Escherichia coli, Enterococcus faecalis*, and *Pseudomonas aeruginosa*, sub-MIC of antibiotics with different modes of action and structures were reported to induce the emergence of biofilm (Kaplan, 2011; Song et al., 2016). Many previous reports proposed that treatment with the sub-inhibitory concentration of various antibiotics set off biofilm formation in *P. aeruginosa* PAO1. Aminoglycosides were previously found to initiate biofilm formation in *P. aeruginosa* isolates (Hoffman et al., 2005). Research conducted on antibiotic resistance suggests that about 65% of hospital-acquired infections are caused due to bacterial biofilms (Potera, 1999). Previous studies report that in presence of antibiotics, the bacterial cells exhibit one or more resistance mechanism(s) to withstand the assault with alterations at the phenotypic level, *i*.*e*., the potential to produce biofilm, which is an adaptive type of resistance. Through this present study, the potency of gentamicin is determined at the sub-minimum inhibitory concentrations on biofilm formation against the environmental strains of *P. aeruginosa* KPW.1-S1 and HRW.1-S3 grown in BH-2% glucose. The present study’s objective was to know how gentamicin at the sub-MIC levels triggers biofilm formation and affects the arrangement of bacterial cells in the biofilm matrix.

Firstly, the biofilm-forming capability of the *Pseudomonas aeruginosa* KPW.1-S1 and HRW1.S3 strains was evaluated by a 24-well microtiter assay method in the presence of 0.75 µg/ml and 1.5 µg/ml of gentamicin which showed increased biofilm load (Figure 2a and 2c). Gentamicin treatment results in biofilm matrix modifications mainly by enhancing eDNA and exoproteins abundance levels, which also favors forming the 3D structures. eDNA has been found to play a structural as well as pivotal role in biofilm origination, binding and shielding biofilms from aminoglycosides, and inducing antimicrobial peptide resistance mechanisms (Chiang et al., 2013; Lewenza et al., 2013; Mulcahy et al., 2008). The structure of the biofilm matrix is composed of extracellular polysaccharides, extracellular DNA, exolipids, alongwith exoproteins. Extracellular DNA (eDNA) was termed as leftovers from lysed cells until Mattick and coworkers discovered that bacterial cells actively release the DNA outside their cells. Exopolysaccharides may be synthesized extracellularly or synthesized within the cells and secreted into the outside environment of their cells (Nwodo et al., 2012). SEM data showed the biofilms adhere to cell surfaces as linear or branched strands that protrude out to form large networks and such scaffolding may be formed with the aid of exopolysaccharides. Exopolysaccharides also serve as scaffolds for other nucleic acids, proteins, carbohydrates, and lipids to adhere. The structures, components, and properties of the exopolysaccharides vary from one to another (Bales et al., 2013). Nelson and co-workers studied the compositions and linkage pattern of EPS matrix from *P. aeruginosa, E. faecalis, K. pneumonia, S. aureus, A. baumanii*, and *E. spp*. biofilms (Bales et al., 2013). Galactose, mannose, and glucose are abundant carbohydrates, followed by N-acetyl-glucosamine, galacturonic acids, arabinose, rhamnose, fucose, and xylose. Exopolysaccharides are not primarily specific to biofilm, but an abundance in their quantity has been reported to be increased thereby resulting in stress responses, such as alginate synthesis in *P. aeruginosa* and colonic acid production in *Escherichia coli* (Laverty et al., 2014). Three exopolysaccharides are associated with *P. aeruginosa* biofilms Pel, Psl, and alginate. Alginate is made up of D-mannuronic acid residues interspersed with L-guluronic acid residues. Alginate confers resistance to *P. aeruginosa* cells from some antibiotics such as ciprofloxacin, ticarcillin, ceftazidime, and gentamicin and it suppresses the immune response of the host (H.-C. Flemming& Wingender, 2010). Although there may be a contribution of exopolysaccharides in biofilm formation by *Pseudomonas aeruginosa* KPW.1-S1 and HRW.1-S3, it is apparent that the presence of antibiotics could not induce more abundance of exopolysaccharides in the case of KPW.1-S1 and HRW.1-S3 biofilms.

The intrinsic characteristics of these biofilms were further highlighted by the topological feat ures of the biofilm architecture and the biofilm assay results correlated with the average biomass and mean thickness of biofilms calculated from confocal imaging. In presence of 0.75 µg/ml of gentamicin, the thickness of biofilm, biovolume, and kurtosis value was increased while skewness was found to be decreased across all concentrations for KPW.1-S1 and HRW.1-S3. The result established that the multicellular assemblage was markedly increased in the presence of 0.75 µg/ml and 1.5 µg/ml of gentamicin. The lesser skewness value of the biofilms of KPW.1-S1 and HRW.1-S3 in the presence of 0.75 µg/ml and 1.5 µg/ml of gentamicin means the biofilms had more void spaces. The greater value of standard deviation correlates to more heterogeneous biofilms of the KPW.1-S1 and HRW.1-S3 strains. It has been established by multiple studies that sub-MIC of a few antibiotics can reduce biofilm production but they cannot kill the bacteria. For example, azithromycin, at low concentrations effectively prevents the formation of *P. aeruginosa* biofilms (Kumar et al., 2021; Zegans et al., 2009). Contrarily, multiple reports demonstrated that several antibiotics at low doses can considerably promote the formation of biofilm in different kinds of bacterial species (Kaplan, 2011; Penesyan et al., 2020).

In previous studies, eDNA was shown to carry out a major structural part in the formation of biofilm by binding and shielding biofilms from the aminoglycosides and inducing resistance mechanisms by antimicrobial peptides in *Pseudomonas aeruginosa* (Tahrioui et al., 2019). Sub-MIC of vancomycin in biofilms of *Staphylococcus epidermidis* (Doroshenko et al., 2014) and sub-MIC of tobramycin in *Pseudomonas aeruginosa* induce eDNA (Tahrioui et al., 2019). In the present study, gentamicin and tetracycline at their Sub-MIC conditions showed a significantly enhanced presence of eDNA in the EPS of *Pseudomonas aeruginosa* biofilms. In the present study, the abundance of exoproteins in EPS was found to be increased significantly in the case of both strains of *Pseudomonas aeruginosa*. A previous report also showed that exoproteins of *K. pneumoniae* biofilm significantly increased in the presence of levofloxacin (Zhang et al., 2021). The Sub-MIC of clindamycin and azithromycin were reported to induce exoprotein production in *S. aureus* (Hu et al., 2019).

Exolipids are also found to be usually predominant in the bacterial biofilm matrix (Flemming and Wingender, 2010). Biochemical analysis depicted that usually, the total dry weight of the extracellular matrix constitutes 15% lipids along with 25% carbohydrates, 55% glycoprotein, and 5% nucleic acids (Alim et al., 2018). In the present study, the presence of exolipids was evident in the EPS of *Pseudomonas aeruginosa* biofilms. For the strain KPW.1-S1, the exolipid content was found to be increased in presence of antibiotics. However, for the strain HRW.1-S3 strain, the abundance of exolipid in biofilms was found to be not altered in presence of antibiotics.

Reactive Oxygen species showed a positive correlation with an increase in the concentration of gentamicin showing an oxidative burst. The oxidative burst damages DNA due to the generation of free radicals causing cell death which may be the reason for an increase in eDNA.

The cell surface hydrophobicity also increased and shifted from hydrophilic to hydrophobic phase showing more adhesive property of the strain resulting in better attachment and biofilm formation. It was observed that ROS generation also increased in the presence of gentamicin. The ROS generated may be due to the oxidative stress prooxidant expression of defense regulons that act with the direct scavenging of active oxygen species (Smirnova et al., 2009). In conclusion, the results obtained from the present study are clinically important as these results indicate that bacterial infections subjected to low doses of antibiotics at the starting and the end of therapy or in the case of treatment with continually low doses of antibiotics may trigger bacterial biofilm formation and thereby may confer resistance to antibiotic treatment and/or emergence of resistant strains. It can be concluded that for *P. aeruginosa* (KPW.1-S1 and HRW.1-S3 strains) the increased biofilm loads in the presence of sub-MIC of gentamicin were mainly associated with the increased abundance of eDNA and exoprotein in EPS structure. Moreover, it was observed that cell surface hydrophobicity increased with increasing concentration of gentamicin depicting an increase in adherent property mediated by cell surface hydrophobicity.

The result of this present study thus provides a scope for future research on antibiotic-mediated increased or decreased biofilm formation in the case of pathogenic bacteria like *Pseudomonas aeruginosa* which is one of the major reasons of chronic diseases.

## Acknowledgments

All authors would like to acknowledge Mr. Ritabrata Ghosh and Mr. Kashinath Sahu for their technical support of confocal laser scanning and scanning electron microscopy respectively.

## Author Contributions

AK and TKS conceived the study and designed the experiments. AK performed most of the experiments. SKS performed experiments on ROS generation and cell surface hydrophobicity. AK, SKS, PB, and TKS analyzed the data and wrote the manuscript.

## Availability of Data and Materials

In this study, all data generated or analyzed are included in the article.

## Ethical Approval

Not applicable.

## Consent to Participate

All authors had consented to participate in the study.

## Consent for Publication

All authors have given consent for publication.

